# Phenotypic diversity acts as a higher-order functional trait of populations

**DOI:** 10.64898/2026.09.24.754102

**Authors:** Kotone Nishikawa, Daiki X. Sato, Yuma Takahashi

## Abstract

Functional traits are usually assigned to individuals, yet some properties exist only as differences among population members. Whether such higher-order properties predict collective performance remains unclear. Here we used the aquatic plant *Spirodela polyrhiza* to test whether phenotypic and genetic differences among genotypes predict non-additive population growth. We grew twelve Japanese strains alone and in all pairwise mixtures and quantified their growth dynamics. Mixtures sometimes outperformed the mean of their component monocultures, and some exceeded the better-performing monoculture, although diversity effects were negative on average. Positive effects were most strongly associated with between-strain differences in frond colour and morphology, identifying specific dimensions of phenotypic diversity as population-level predictors of synergistic growth. Genome-wide higher-level association mapping identified loci where between-strain diversity was associated with growth effects, implicating stress responses, secondary metabolism and frond morphology. Loci associated with reduced performance implicated chemical perception and self/non-self recognition, whereas those associated with positive effects reflected functional differentiation in stress responses and physiological niches. Our findings suggest that some traits generate new functions at the population level through variation among individuals, making diversity itself a higher-order functional property.

## Introduction

Biological function is commonly attributed to traits of individuals. Yet some properties of populations are defined not by the phenotype of any single individual, but by differences among their members. Such higher-order properties may affect collective performance, but their phenotypic and genetic bases remain poorly understood. Changes in collective performance caused by variation among constituent elements are broadly referred to as diversity effects. In species-rich communities, diversity effects are often explained by selection and complementarity effects, whereby variation among species enhances collective performance through dominance by high-performing taxa, resource partitioning, facilitation, or insurance against environmental fluctuations ^1–5^. Whether analogous principles explain the ecological consequences of diversity within populations remains less clear. Intraspecific diversity differs from species diversity because individuals often differ along continuous genetic and phenotypic axes, interact with close relatives or non-relatives, and generate population-level outcomes that depend on the particular combinations of genotypes present in a population ^6–10^.

Genetic and phenotypic variation within populations can enhance group performance, productivity, resistance to disturbance, and stability ^7,11–20^. However, such effects are not universal. Within-population diversity can have weak, neutral, or even negative effects on population performance, and these effects can arise non-additively from interactions among genotypes ^17,19–23^. Thus, a central challenge is not simply to ask whether diversity is beneficial, but to identify which dimensions of genetic and phenotypic variation determine the magnitude and direction of diversity effects. Unlike conventional individual traits, these dimensions are defined by differences among the members of a population and therefore represent candidate higher-order functional properties.

Addressing this challenge requires shifting the unit of association from individuals to populations. Conventional genome-wide association studies (GWAS) are designed primarily to associate allelic variation in individual genomes with individual-level phenotypes ^24,25^. They cannot directly identify properties whose values are defined only by differences among individuals, such as phenotypic or locus-specific diversity within a population. This distinction is critical because population dynamics in genetically heterogeneous groups can depend not only on the intrinsic performance of each genotype, but also on interactions among genotypes, including indirect genetic effects and other social or ecological effects mediated by neighboring individuals ^26–28^. A framework is therefore needed in which diversity itself, measured at each genetic locus or phenotypic trait, is used to explain non-additive population growth. To address this gap, we developed genome-wide higher-level association study (GHAS) and phenome-wide higher-level association study (PheHAS) as methodological frameworks for identifying the higher-order genetic and phenotypic bases of diversity effects ^18,20^. GHAS identifies genetic loci at which within-population genetic diversity explains variation in diversity effects, whereas PheHAS identifies phenotypic traits at which within-population phenotypic diversity explains such variation. Identifying these higher-order bases can clarify the ecological mechanisms underlying diversity effects, reveal the evolutionary processes that generate and maintain useful diversity, and guide the design or assembly of populations that elicit beneficial collective outcomes.

Here, we apply this framework to the duckweed *Spirodela polyrhiza*, a free-floating clonal aquatic plant. Duckweeds provide a tractable system for studying diversity effects because genetically defined lineages can be propagated rapidly, assembled into controlled mixtures, and phenotyped under standardized conditions ^29,30^. *Spirodela polyrhiza* is also suitable for genomic and transcriptomic analyses because its genome and transcriptome have been characterized, and it has one of the smallest known monocot genomes ^31,32^. Using this system, we identify specific phenotypic and genetic differences among genotypes that predict non-additive population growth, providing a test of whether population-level properties defined by variation among individuals can be mapped empirically. First, we quantified population growth across 12 monocultures and all 66 pairwise genotype mixtures, using time-series imaging to estimate intrinsic growth rate, carrying capacity and overall growth dynamics. Second, we quantified synergistic growth by estimating the diversity effect and transgressive effect for each genotype combination. Third, we tested which differences in frond morphology, colour, root length and growth characteristics among genotypes predicted these synergistic effects. Finally, using RNA-seq-derived genetic variants and genome-wide higher-level association analyses, we identified genomic regions and candidate genes associated with variation in synergistic growth.

## Results

### Phenotypes and population parameters

The variation in frond area, root length, frond shape and the three colour parameters, L* (lightness), a* (the axis from green to red) and b* (the axis from blue to yellow), varied significantly among strains (Fig. 2A, Supplemental Information Fig. S1). Correlations among traits were generally weak (Fig. S2); however, L* and b* were highly collinear and exhibited nearly identical variation. Consequently, L* was excluded from subsequent analyses. Population growth was quantified from time-series images as total frond-cover area per container. Logistic growth models fitted to these trajectories provided estimates of intrinsic growth rate (*r*) and carrying capacity (*K*). Across nearly all experimental treatments, population growth reached a plateau by day 20 (Fig. 2B). In the principal component analysis, population sizes at all time points exhibited loadings in the same direction on the first principal component (PC1), indicating that it represents overall growth capacity (Fig. 2C). PC1 explained 69.3% of the total variance. Note that the higher values of PC1 indicate a lower growth ability. For better interpretability, the value of PC1 multiplied by −1 was used in the subsequent analyses as an index of growth ability. In monoculture treatment, *r* showed no significant difference among the 12 strains (χ² = 15.2, *P* = 0.17) but *K* significantly varied among strains (χ² = 26.0, *P* = 0.006) (Fig. 2D). PC1 differed significantly among the 12 strains (χ² = 44.1, *P* < 0.001). On the other hand, in polyculture treatment, there were significant difference among the strain combinations for *r* (χ² = 139.2, *P* < 0.001), *K* (χ² = 101.1, *P* = 0.001), and PC1 (χ² = 181.9, *P* < 0.001) (Fig. 2E). Under monoculture conditions, a significant correlation was detected only between *r* and PC1 (Fig. 2F), whereas under polyculture conditions, significant correlations were observed among all combinations of parameters (*r*, *K*, and PC1) (Fig. 2G).

**Figure 1.**
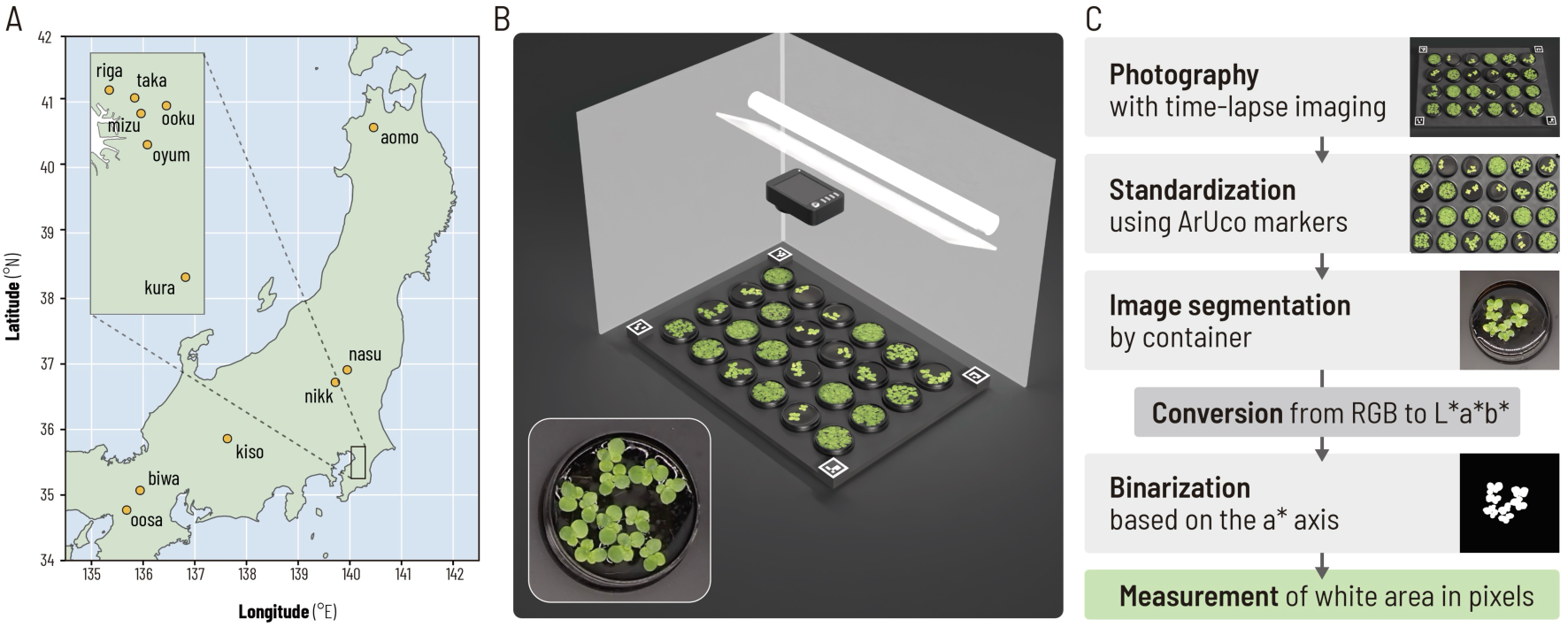
Sampling locations and overview of the experimental and image-analysis procedures. A) Sampling locations. Sampling sites were distributed across Japan, spanning approximately 35°N to 41°N. B) Imaging setup. Plants were cultured and photographed using time-lapse imaging inside a booth surrounded by white walls. LED illumination was provided from above through a diffuser. ArUco markers placed around the culture plate were used for image standardization. The inset shows an example of a culture container during the experiment. C) Image-analysis workflow. Images were standardized using ArUco markers, segmented by culture container, converted from RGB to L*a*b* color space, and binarized based on the a* axis. Plant area was quantified as the number of white pixels in the binarized image.

**Figure 2.**
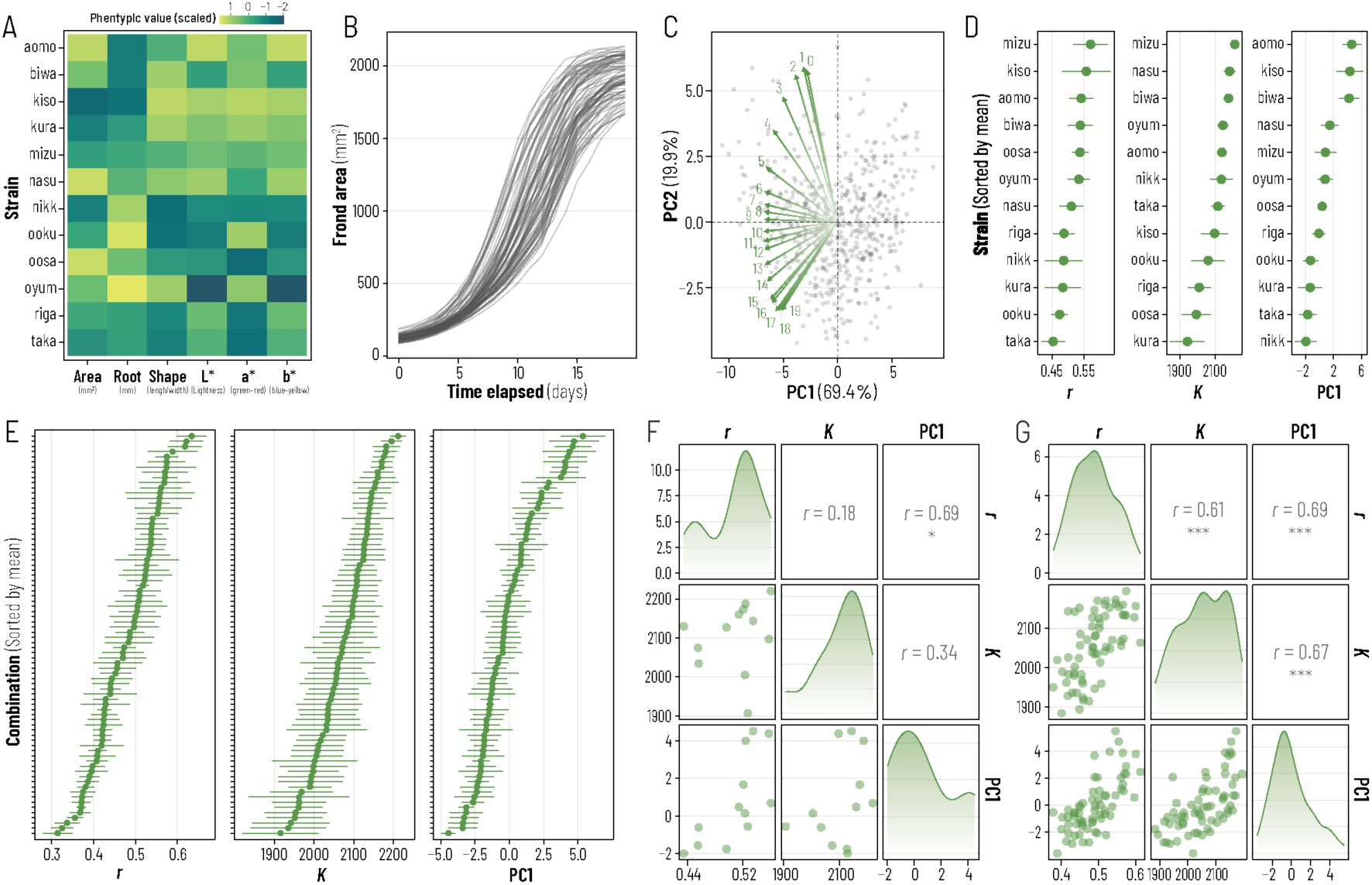
Phenotypic traits and growth characteristics of the duckweed strains. A) Heatmap showing frond area, root length, frond shape (length-to-width ratio), frond lightness (L*), a* value (green–red axis), and b* value (blue–yellow axis) for each strain. Values were scaled for visualization. B) Changes in frond area over the culture period. Frond area increased over time and generally approached a plateau after 20 days. Each line represents one strain combination. Insets show representative images recorded at different stages of growth. C) Principal component analysis of temporal growth patterns. PC1 primarily represented overall growth rate and frond area and, in its original orientation, larger PC1 scores corresponded to lower growth performance. Therefore, PC1 scores were multiplied by −1 for subsequent analyses so that larger values represented greater growth performance. PC2 represented variation in the timing of growth, with higher PC2 scores indicating growth patterns in which frond area was relatively larger during the earlier part of the culture period. Numbers indicate the time points used in the analysis. D) Intrinsic rate of increase (*r*), carrying capacity (*K*), and PC1 of growth patterns for each strain grown in monoculture. Points indicate mean values and horizontal bars indicate ± SE. E) *r*, *K*, and PC1 for each strain combination grown in polyculture. Points indicate mean values and horizontal bars indicate ± SE. F) Pairwise correlations among the three growth parameters (*r*, *K*, and PC1) in monocultures. G) Pairwise correlations among the three growth parameters in polycultures.

### Overyielding and transgressive overyielding

We quantified diversity effects as the difference between each mixture’s observed growth parameter and the mean value of its two component monocultures. Observed values deviated significantly from these monoculture-based expectations (Fig. 3A–C). Wilcoxon signed rank tests also confirmed significant differences between the observed and expected values for *r* (*V* = 720, *P* = 0.013), *K* (*V* = 559, *P* < 0.001) and PC1 (*V* = 692, *P* = 0.008). For *r*, *K* and PC1, the observed values were higher than the expected values in 25, 24 and 25 of the 66 combinations, respectively. Thus, combinations exhibiting overyielding accounted for approximately 37% of all cases across the population parameters examined, and the average magnitude of overyielding was biased toward negative values (left panels of Fig. 3D–F). Moreover, combinations showing transgressive overyielding (i.e., exceeding the higher value of the two component monocultures) for *r*, *K* and PC1 accounted for 27%, 17% and 17% of the combinations, respectively (right panels of Fig. 3D–F).

**Figure 3.**
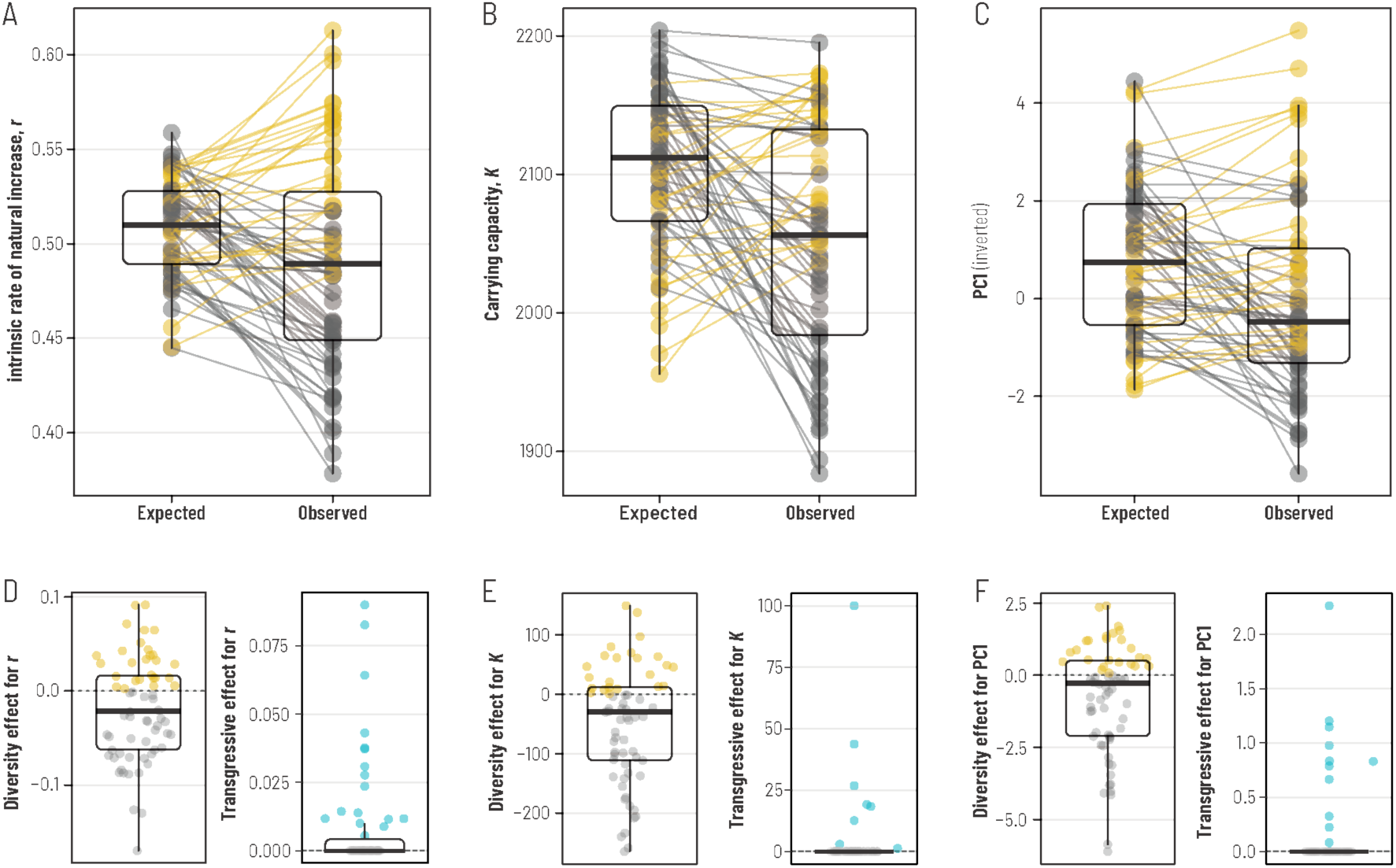
Expected and observed growth characteristics of polycultures and their diversity/transgressive effects. A–C) Expected and observed values of the intrinsic rate of increase (*r*; A), carrying capacity (*K*; B), and PC1 of growth patterns (C) in polycultures. Expected values were calculated from the corresponding monocultures of the two strains constituting each polyculture, whereas observed values were obtained from the actual two-strain mixtures. Paired points connected by lines represent individual strain combinations. D–F) Diversity effects and transgressive overyielding effects for *r* (D), *K* (E), and PC1 (F).

### Phenotypic basis of overyielding

We next tested whether between-strain differences in phenotypic traits and monoculture growth parameters predicted diversity effects. Multiple regression models included standardized pairwise distances and their second-order interactions, with genetic distance included as a covariate. Because significant interaction terms were detected, main effects should be interpreted with caution; accordingly, when interaction terms were retained in the selected models, we did not further discuss the corresponding main effects of the variables involved. Model selection for the regression analysis of overyielding in *r* indicated that interaction terms involving *r*, b*, and area were included in the best-supported model and exhibited relatively large positive regression coefficients (Fig. 4A). In the model selection for *K*, interaction terms including *r*, root length, and shape were identified as having relatively large positive coefficients (Fig. 4B). Similarly, model selection for PC1 revealed that the interaction terms involving *r*, root length, shape and area showed relatively large positive regression coefficients (Fig. 4C). The interaction effect between *r* and shape was consistently detected as a strong negative effect across all dependent variables.

**Figure 4.**
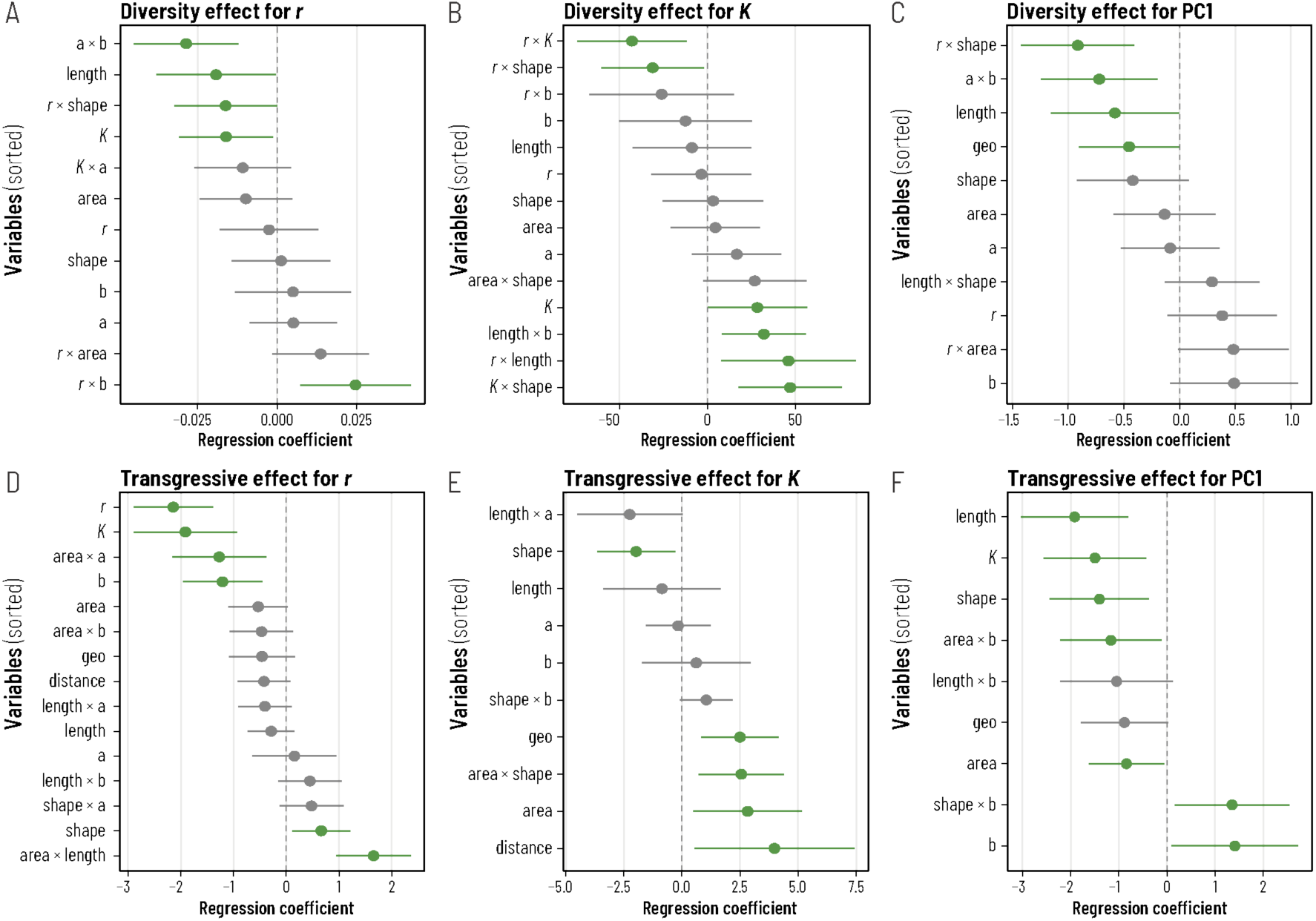
Multiple regression analyses of factors associated with diversity and transgressive effects on growth parameters. Forest plots show regression coefficients from the best-supported models, with diversity effects (A–C) or transgressive effects (D–F) as the response variables. A, D) Intrinsic rate of increase (*r*). B, E) Carrying capacity (*K*). C, F) PC1 of growth patterns. Points indicate estimated regression coefficients, and horizontal bars indicate ± SE. Green points and lines indicate statistically significant coefficients. When an interaction term was significant, the corresponding main effects were retained in the model but were not interpreted independently.

For the transgressive effect on *r*, variables such as shape and the interaction between area and root length were included in the best-supported model as predictors with relatively large regression coefficients (Fig. 4D). For the transgressive effect on *K*, the main effect of genetic distance and the interaction effect between area and shape showed a relatively large effect (Fig. 4E). Similarly, for the transgressive effect on PC1, interaction effect of b* and shape were selected as variables with relatively large positive effects (Fig. 4F).

### Transcriptome-based association analysis

Using the full set of genes analysed, we found no significant pattern of isolation by distance (i.e., increasing genetic distance with greater geographic distance) among the strains in this population (Fig. S3). To identify genomic regions associated with diversity effects, we performed GHAS using pairwise nucleotide diversity (π) within non-overlapping 5-kb windows. For each window, π was tested for association with diversity effect and transgressive effect across the 66 strain pairs, accounting for background genetic similarity (Fig. S4). Among the top 50 loci showing significant associations, the number of detected genes differed substantially among growth metrics (*r*, *K* and PC1), with limited overlap at the gene level (Fig. 5A). Nevertheless, a small subset of genes was shared between pairs of metrics, particularly between *K* and PC1 or between *r* and PC1. We then conducted gene ontology (GO) enrichment analysis using the list of the top-associated genes (Fig. 5B). GO enrichment analysis identified distinct functional associations with diversity effects. Greater nucleotide diversity in genes involved in hydroxycinnamoyl transfer and the positive regulation of flavonoid biosynthesis was associated with stronger positive transgressive overyielding effects on *K*. In contrast, greater diversity in genes associated with phospholipid catabolism, fatty-acid and eicosanoid transport and secretion, unsaturated-fatty-acid biosynthesis, methionine–homocysteine metabolism, and somatic cell DNA recombination tended to reduce transgressive overyielding, shifting both *K* and the PC1 score of growth patterns toward the absence of overyielding. Among the loci associated with overyielding and transgressive overyielding, one of the strongest signals of association with transgressive overyielding in *K* was detected on chromosome 4, corresponding to a non-synonymous substitution (E592G) in the gene Sp9512.a02.4.g05610 (Fig. 5C). This variant was predicted by PROVEAN to affect protein function (Provean score = −2.79; Table S3) and showed significant differences in frond shape (*P* = 0.020, Wilcoxon’s rank-sum test; Fig. 5D), indicating that the E592G substitution is linked to morphological variation at the individual level. Moreover, genotype pairs differing at this position exhibited higher transgressive overyielding than pairs carrying the same allele (*P* = 0.049, Wilcoxon’s rank-sum test; Fig. 5E). Based on the AlphaFold-predicted structure, the E592 residue is located within a pentatricopeptide repeat (PPR) motif and lies at a position likely to influence the local repeat conformation (Fig. 5F).

**Figure 5.**
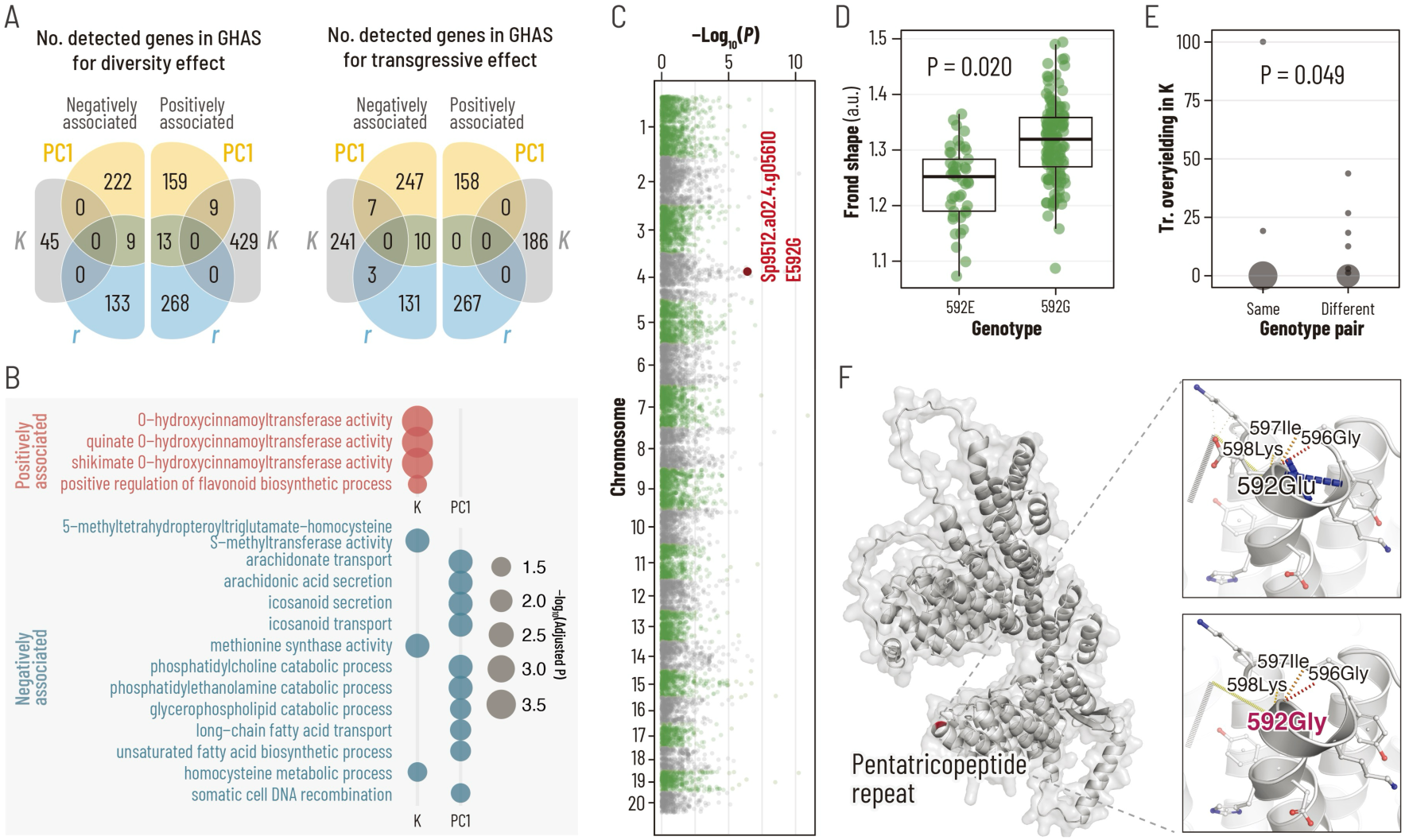
Genomic associations with diversity-driven growth effects. A) Overlap among genes located in the top 50 genomic windows associated with diversity effect and transgressive effect for growth rate (*r*), carrying capacity (*K*), and PC1. For each metric, genes in negatively associated windows are shown on the left and those in positively associated windows on the right. B) Gene ontology terms enriched among the top-associated genes. Point color indicates the direction of association (red, positive; blue, negative), and size represents −log_10_(*P*). C) Genome-wide association profile for transgressive overyielding in *K*. The locus containing the nonsynonymous E592G substitution in Sp9512.a02.4.g05610 is highlighted. D) Frond-shape traits of strains carrying the E592 or G592 allele. E) Transgressive overyielding in *K* for strain pairs carrying the same or different alleles at residue 592. F) AlphaFold-predicted structure of the protein encoded by Sp9512.a02.4.g05610, with enlarged views of the E592 and G592 variants within a pentatricopeptide repeat motif.

## Discussion

Our results reframe a long-standing question in biodiversity research by shifting attention from the traits of individuals to the differences among them. Rather than asking only whether genetic diversity benefits a population, we ask which phenotypic and genetic differences among individuals predict collective performance. These differences constitute candidate higher-order functional properties because their values are defined at the level of genotype combinations rather than single individuals. Classical explanations of diversity effects invoke complementarity and selection as general mechanisms ^1,3,4,33^, and within populations, genetic variation has repeatedly been shown to enhance productivity and stability ^7,11,13,34,35^. In *Spirodela polyrhiza*, however, overyielding and transgressive overyielding varied widely among genotype combinations and were, on average, negative. Genetic divergence alone was a poor predictor of collective performance, and most mixtures underperformed their monoculture expectation, consistent with reports that intraspecific diversity can be neutral or even detrimental ^17,21,22^. Yet a subset of combinations grew synergistically, and these were not randomly distributed. Phenotype-level (PheHAS) and genome-level (GHAS) higher-order association analyses ^18,20^ converged on a restricted set of functional dimensions: frond colour, shape and size at the phenotypic level, and stress response, secondary metabolism and frond morphology at the genetic level. Diversity effects were therefore associated not with divergence per se, but with divergence along specific phenotypic and genetic axes. In this sense, phenotypic diversity is not merely a measure of heterogeneity: when defined along a functionally relevant axis, it becomes a population-level property associated with collective performance. This distinction between diversity as a quantity and diversity as a multidimensional composition provides a bridge from variation among genotypes to emergent population growth.

Which dimensions of phenotypic variation, then, generate synergy? Variation in frond colour emerged as one of the strongest predictors of enhanced diversity effects, with consistent patterns identified by multiple regression several performance metrics. Leaf colour can reflect variation in chlorophyll abundance, photosynthetic capacity and pigment composition, as well as physiological responses to environmental stress ^36–38^. This physiological interpretation is partly supported by our genomic analyses, which identified enriched functions related to stress-responsive pigment and phenylpropanoid metabolism, including positive regulation of flavonoid biosynthetic process and O-hydroxycinnamoyltransferase activity. Flavonoids contribute to antioxidant protection and can reduce oxidative damage in photosynthetic tissues, whereas hydroxycinnamoyltransferases occupy a central position in phenylpropanoid metabolism and influence the production of structurally and physiologically diverse phenolic compounds ^39,40^. Although these genomic signals do not establish a direct genetic basis for frond colour variation, they indicate that differences among strains in pigment-associated metabolism and protection against physiological stress may underlie the observed relationship between colour divergence and diversity effects. Variation along the blue-to-yellow axis (b*) was especially informative. Leaf yellowing commonly accompanies chlorophyll degradation and the loss or reorganization of photosynthetic proteins during senescence and abiotic stress ^38,41^. Differences in b* may therefore capture biologically meaningful variation in physiological state, environmental optima or stress tolerance among genotypes. Culture environments are inherently dynamic, with resource availability, physicochemical conditions and population density changing over time. Under such conditions, combining genotypes that differ in pigment metabolism, antioxidant capacity or stress-response properties may broaden the range of conditions under which population growth can be sustained. Such temporal or physiological complementarity could enhance non-additive population performance in a manner consistent with niche complementarity. Frond-colour differences may therefore indicate a broader axis of physiological and stress-response differentiation among genotypes. Diversity along this axis is a candidate higher-order functional property that could promote complementarity, although the underlying physiological differences were not measured directly here.

A second functional axis may operate through an entirely different mechanism. Frond morphology and size were also identified as important determinants of diversity effects, and our genomic analyses recovered loci associated with frond morphology, suggesting that these morphological differences have a genetic basis. In *S. polyrhiza*, the upper and lower frond surfaces have distinct structures which, together with variation in frond curvature, thickness and ventral shape, may influence local deformation of the air–water interface. Such deformations can generate capillary attraction or repulsion between floating bodies and interactions with container walls, a phenomenon commonly termed the Cheerios effect ^42–44^. Frond morphology may therefore influence not only structural properties but also spatial behaviour within populations. Consistent with this possibility, genotype combinations with greater differences in frond size also showed greater differences in their tendency to associate with the container wall (Fig. S5). When genotypes with contrasting interfacial properties are mixed, such as those predominantly associated with upward versus downward surface deformation, their interactions with the water surface at the container wall may cause them to preferentially occupy different regions, for example, the center versus the margin. This spatial segregation could redistribute individuals and reduce local crowding. Because high population density suppresses duckweed growth ^45,46^, such spatial reorganization may alleviate density-dependent suppression and thereby contribute to positive diversity effects. Thus, morphological differences among genotypes may generate a population-level property, i.e., spatial organization, that no individual frond possesses alone. Morphological diversity could thereby affect collective performance through capillary interactions and the redistribution of individuals across the water surface.

The functional axes identified above account for the synergistic minority, but they leave the majority unexplained. In our system, diversity effects were predominantly negative, and positive overyielding and transgressive overyielding occurred in only a limited subset of genotype combinations. This asymmetry suggests that beneficial diversity effects arise only when particular forms of functionally complementary variation are present, whereas many combinations instead generate neutral or suppressive interactions. Negative diversity effects can emerge when niche overlap intensifies competition or when strongly competitive genotypes suppress their neighbours, offsetting any benefits of complementarity (Loreau and Hector 2001; Schmid et al. 2008). Other non-mutually exclusive mechanisms may include physiological incompatibility, differential sensitivity to conditions created by neighbouring strains, and chemically mediated responses to non-self genotypes. Experimental studies in terrestrial plants have shown that soluble root exudates allow plants to distinguish self, kin and unrelated neighbours and can induce changes in root proliferation and competitive allocation ^47,48^. Although equivalent recognition mechanisms have not yet been demonstrated in *S. polyrhiza*, continuously shared culture media provide a plausible route through which genotype-specific metabolites could alter the growth of neighbouring fronds. Because our mixtures included strains spanning substantial genetic distances, some genetically divergent combinations may have elicited stronger mutual or asymmetric responses, but this possibility remains to be tested directly. The predominance of negative interactions does not weaken the utility of our framework; rather, it emphasizes that both tails of the diversity-effect distribution are biologically informative. Linking phenotypic and genetic differences to population performance can identify not only the traits and loci associated with synergistic growth, as demonstrated by genetic analyses of overyielding in other genotype mixtures, but also those associated with competitive dominance or suppressive interactions ^49^. A framework that considered only beneficial outcomes would therefore misrepresent how intraspecific diversity operates in populations.

The enriched functions described below identify candidate mechanisms rather than demonstrate them, because divergence in a gene set does not establish that it mediates the interaction. One candidate mechanism underlying these negative effects is chemically mediated interaction cost among genetically distant strains. Genetic divergence in genes associated with phospholipid catabolism, unsaturated-fatty-acid biosynthesis, and lipid transport or secretion may produce genotype-specific chemical environments in the shared water column. Plants can alter competitive traits in response to chemical cues from kin and non-kin neighbours ^47,50–53^, suggesting that chemically dissimilar strains could induce stronger competitive or stress responses. The enrichment of phosphatidylcholine and glycerophospholipid catabolism supports a role for membrane remodelling and lipid-derived metabolites, whereas long-chain fatty-acid transport and lipid-secretion terms are consistent with their movement into the extracellular environment, where lipid-derived compounds can participate in plant stress signalling ^54–56^. Arachidonate- and icosanoid-related GO terms should be interpreted cautiously because canonical arachidonic-acid-derived eicosanoid signalling is primarily characterized in animals, whereas lipid-derived signalling in higher plants is dominated by C18 fatty-acid-derived oxylipins ^55^. These terms may therefore reflect broader functions in lipid transport and secretion rather than an animal-like eicosanoid pathway in duckweed. Enrichment of methionine synthase activity and homocysteine metabolism further suggests that divergence in one-carbon and sulfur metabolism may contribute to metabolic or stress-related interaction costs. Together, these functions are consistent with the possibility that genetically distant strains generate dissimilar chemical environments, although they do not identify the compounds or recognition pathways involved. Conditioned-medium and exudate-transfer experiments could test whether growth suppression is mediated by diffusible compounds in the absence of physical contact.

The two mechanisms identified here, physiological complementarity and morphology-dependent spatial restructuring, differ in almost every respect, yet both were recovered by the same analytical logic. This points to a more general methodological argument. Conventional genome-wide and phenome-wide association studies are designed to link the genotype or phenotype of an individual to that individual’s own traits ^24,25^. Such designs cannot, in principle, test whether diversity at a locus explains a property that no single individual possesses. Population growth in a genetically heterogeneous group is precisely such a property: it depends not only on the intrinsic performance of each genotype but on interactions among them, including indirect genetic effects and other social or ecological effects mediated by neighbours ^26–28^. GHAS and PheHAS resolve this mismatch by making diversity itself, measured at each locus or trait, the explanatory variable, and emergent non-additive performance the response ^18,20^. The unit of association shifts from the individual to the population, and the question shifts from which alleles make an individual perform well to which allelic and phenotypic differences make a collective perform well. In this framework, diversity along a particular phenotypic or genetic axis can be treated as a higher-order functional property when it predicts performance emerging from interactions among population members.

The generality of this approach does not depend on duckweed. It requires only that genetically defined lineages of a single species can be assembled into replicated mixtures and that a collective performance metric can be measured. These conditions are met in microbial strain panels, crop cultivar mixtures, and systems in which intraspecific diversity effects were first demonstrated, including eelgrass genotype mixtures ^7,11^ and *Solidago* genotype assemblages ^13^. Although these studies established that genotypic richness can enhance productivity and resistance, they were not designed to identify the components of genetic variation responsible for these effects. Higher-order association mapping provides a means to connect genomic and phenotypic variation with emergent group performance, extending quantitative and community genetics beyond individual phenotypes ^8,57,58^. It may also reveal conflicts between individual and collective performance. For example, if chemically mediated non-self responses suppress genetically dissimilar competitors, behaviour advantageous to an individual may reduce the productivity of the group it inhabits. The genetic architecture of competitive success may therefore differ from that of collective performance, a tension that higher-order association mapping is well positioned to expose. Its recovery here of physiological, morphological and biophysical axes, together with a candidate recognition-mediated cost of divergence, illustrates its capacity to reveal mechanisms not apparent from individual-level analyses.

Allelic differences that generate negative diversity effects may be subject to positive frequency-dependent selection, which tends to eliminate variation within populations by favouring locally common alleles ^15^. Our mixtures, however, combined strains collected from geographically distinct populations separated by up to 800 km. Alleles fixed or maintained independently in different populations can therefore be diverse across our strain panel even if they do not normally coexist within local populations. Bringing these divergent lineages together may expose antagonistic interactions or incompatibilities that are rarely expressed in nature, making the predominance of negative diversity effects unsurprising. Our results thus characterize the genetic and phenotypic consequences of combining geographically differentiated lineages under defined conditions; whether the same loci vary within natural populations and influence their collective performance remains to be tested.

Intraspecific variation can have ecological effects comparable in magnitude to differences among species ^10^, yet it is still commonly represented as a single quantity. Our results show that genetic distance alone predicts collective performance poorly and that mixtures of divergent genotypes can perform worse than expected from their monocultures. What distinguishes synergistic combinations is not simply how much variation they contain, but the phenotypic and genetic axes along which their members differ. These results identify differences in frond morphology and colour as population-level predictors of collective growth and point to stress-response differentiation as a candidate genetic axis of complementarity. More generally, they show that functional properties of populations can be defined by differences that no single individual possesses. Diversity should therefore be treated not as a scalar quantity, but as a multidimensional set of higher-order properties whose effects depend on the particular dimensions combined within a population.

## Materials and Methods

### Study species and strains

*Spirodela polyrhiza* is a small, floating aquatic plant characterized by simple, flattened fronds that lack true stems or leaves. Its fronds often float on the water surface, with several thin roots extending downward, facilitating nutrient uptake ^32,59^. They exhibit a rapid clonal propagation strategy primarily through vegetative reproduction: Each frond produces daughter fronds via a meristematic zone. Their populations can double within a few days, reflecting extremely high intrinsic growth rates ^60^. Due to its high growth rate and high starch and amino acid content, it offers significant potential for sustainable food and biofuel production ^61–63^. Its rapid growth and two-dimensional spread on the water surface make it easy to observe population growth using simple image analysis. The absence of sexual reproduction facilitates the evaluation of how genetic heterogeneity influences population dynamics across generations. Additionally, the genome size is remarkably small (180 Mbp), and the entire genome sequence is fully sequenced and accessible (e.g., Wang et al. 2014). Taken together, *S. polyrhiza* serves as an ideal model system for the experimental investigation of the effects of genetic heterogeneity on population dynamics and its genetic basis.

In 2023, *S. polyrhiza* specimens were collected from 12 geographically distinct locations across Japan (Table S1, Fig. 1A). These populations are separated by distances of up to approximately 800 km, while some populations are geographically close to one another. However, all population pairs were separated by more than 2 km and were thus treated as independent populations. Twelve strains were isolated and maintained under laboratory conditions by cultivating a single colony obtained from each site. This species reproduces only asexually; therefore, no genetic variation was observed among individuals within each strain. The strains were maintained in plastic Petri dishes (100 mm in diameter × 40 mm in height) using 100 mL MS medium (Murashige and Skoog medium). Alternatively, NF medium (2.7 mM CaCl_2_, 1.2 mM MgSO_4_, 1 mM KH_2_PO_4_, 5 mM KNO_3_, 18 μM MnCl_2_, 46 μM H_3_BO_3_, 0.77 μM ZnSO_4_, 0.32 μM CuSO_4_, 0.49 μM MoO_3_, 20 μM FeSO_4_, 50 μM Na_2_EDTA) was used, as described by Muranaka et al. (2015). In addition, 100 µg/mL sodium ampicillin, 1 ppm sodium hypochlorite (Milton), or 200 µL/L germanium solution (diatom removal agent, Chagoke Killer) was supplemented. Cultivation was conducted under controlled plant light conditions with a 12-hour or 16-hour photoperiod. Room temperature was kept at 23–25℃. To prevent overgrowth of algae and bacteria, the culture medium was replaced every 2–3 days.

### Phenotypic measurements

Frond size, shape, colour and root length were measured for the 12 strains grown under the conditions described above. Frond area was quantified from photographs taken with a Leica stereomicroscope: RGB images were converted to Lab* colour space in PlantCV (v4.5.1), the a* channel was thresholded at 114, and frond area was calculated from the resulting white pixel count. Root length was measured with a digital caliper, taking the longest root when a frond had more than one. Approximately 15 individuals per strain were measured, using randomly selected parent fronds from different culture containers.

To quantify frond shape and color, fronds were photographed using a digital camera under standardized lighting conditions together with a color reference and a size reference. ImageJ was used to standardize measurements based on the size reference, after which frond width and length were measured for the largest frond (the parent frond) in each colony. Leaf shape was evaluated based on the ratio of length to width. Color calibration was then performed using the color reference in Adobe Photoshop, and mean RGB intensity values were calculated from a 60 × 60 pixels region at the center of the parent frond. RGB values were converted into the CIELAB (L*a*b*) color space. Leaf color expressed in the CIE L*a*b* color space has been widely used to evaluate leaf pigment content and plant health status ^41^. Frond shape and color were measured on approximately 15 individuals per strain.

### Cultivation experiment

Cultivation was performed under monoculture (single strain) and polyculture (two strains) conditions: pairing all 12 strains in every combination gave 12 monocultures and 66 polycultures. Black containers (60 mm diameter × 41 mm height) were filled with NF medium and arranged in a custom 24-well plate (6 × 4). Four fronds of the same strain were introduced per monoculture container, and two fronds of each strain per polyculture container (Fig. 1B). Growth was monitored for 20 days by photographing the plate every 6 h with a digital camera (Olympus Tough TG-6, 4,000 × 3,000 px) positioned 35 cm above the plate; fronds typically covered the water surface in most combinations by the end of this period. Only images from the light period were analysed. Using the ArUco module in OpenCV, four DICT_4X4_50 markers (400 px, one per corner of the plate) served as reference points for a perspective transformation that corrected distortion (Fig. 1C); corrected images were cropped to 2,100 × 1,400 px and split into 24 segments (350 × 350 px), one per container, and only segments in which all four markers were detected were retained. Each segment was converted from RGB to L*a*b* colour space in PlantCV, and the a* channel was extracted, binarized at a threshold of 114, and denoised with a median filter (kernel size 5). Frond cover was quantified as the number of white pixels in the resulting binary image and converted to area for subsequent analyses.

### Population parameter estimation

Before estimating population parameters, we selected and corrected the data for measurement error. Replicates in which total frond cover failed to reach two-thirds of the container’s water surface area over the 20-day cultivation period were excluded, as this growth cessation was primarily due to bacterial proliferation. We also corrected unnatural decreases in frond cover that occurred later in cultivation despite no true reduction in cover; these apparent decreases resulted from failure to detect green colour when fronds became submerged or covered by dark green algae. Where frond cover at a given time step was lower than at the previous step, we replaced it with the previous value, ensuring that frond cover did not decrease over the experimental period in the data used for subsequent analyses.

To extract population growth characteristics for each container, we performed PCA on the time-series frond cover data. Daily average frond cover per container was first calculated from (typically four) data points per day, and PCA was performed on these daily values across all containers.

To estimate population growth parameters, we fitted a logistic model to the time-series data for each container using the *nls* function in *R*, estimating initial population size (*N*_0_), intrinsic growth rate (*r*), and carrying capacity (*K*). When initial growth was slow and population size had not reached *K* within the 20-day cultivation period, *K* tended to be overestimated; in such cases, if the estimated *K* exceeded the container’s water surface area (60,000 pixels, 2,827 mm^2^), it was replaced with 60,000 pixels.

### Calculation of overyielding and transgressive overyielding

The average of population parameters, such as *r*, *K*, and principal components, were calculated for each monoculture strain and each polyculture combination. Overyielding, *OY*, for each polyculture combination was calculated using the following formula:

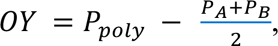

where *_PA_*, *_PB_*, *_Ppoly_* represent the population parameters of monoculture of strain A, monoculture of strain B, and polyculture of these strains, respectively. The first term represents the observed value under polyculture conditions, while the second term corresponds to the expected value derived from the average of two monoculture populations, i.e., null expectation. The difference between these two values represents the non-additive effects generated by interactions among strains, known as diversity effects.

Then, the transgressive over-yielding, *TOY*, in polyculture condition, defined as the increase in yield under polyculture conditions relative to the highest-yielding monoculture condition, was also evaluated using the following formula:

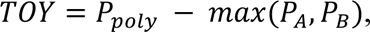

where *P_poly_* represents the observed population parameters for a polyculture consisting of strains A and B. Second term of this formula, *max* (*P*_A_, *P*_B_), represents the larger of the two values: *P*_A_, *P*_B_.

### Phenotype-level association analyses

To identify which aspects of phenotypic diversity explain overyielding and transgressive overyielding, we built multiple regression models with phenotypic distance (in frond area, root length, *r*, *K*, frond shape, a* and b*) and its second-order interactions as explanatory variables, and overyielding or transgressive overyielding for each strain combination as the response variable. Phenotypic distances were calculated as Euclidean metrics, with numerical data standardized before regression. Overyielding was analysed with a Gaussian generalized linear model; because transgressive overyielding contained many zeros, it was analysed with a Tweedie-distributed model using *glmmTMB* (package glmmTMB). Best-fit models were identified by stepwise selection based on the Akaike information criterion (AIC), with genetic distance between cocultured strains included as a correction term.

### RNA-seq analysis and variant calling

A single parental frond of *S. polyrhiza* was harvested from 12 strains at 15:00. RNA was extracted from whole frond with the Maxwell^®^ RSC Plant RNA Kit (Promega, AS1500). Paired-end libraries (150 bp) were sequenced on an Illumina NovaSeq X Plus, which yielded 24–34 million read pairs per sample. Adapter trimming and quality control were performed with fastp v0.23.4 ^65^ using the options -q 30 -u 30 -n 10 -l 20; reads shorter than 20 bp or containing >30% low-quality bases (Q < 30) were discarded. After this QC processing, approximately 93% of reads were retained for downstream analyses (Table S2).

Trimmed reads were aligned to the *S. polyrhiza* reference genome (Sp9512 a02u1, 138 Mb; ^66^) with STAR v2.7.11a ^67^in two-pass mode, retaining uniquely mapped reads only (--outFilterMultimapNmax 1) and allowing up to 4 % mismatches (-- outFilterMismatchNoverLmax 0.04). Alignment files were processed with GATK v4.6.1 following the RNA-seq Best-Practices workflow ^68^: read-group assignment, duplicate marking, SplitNCigarReads, and two rounds of base-quality score recalibration (BQSR). Because validated variant databases are unavailable for *S. polyrhiza*, we generated an internal bootstrap resource from first-round calls and supplied these as known sites for the second round of BQSR.

Variant calling was carried out per sample in GVCF mode using HaplotypeCaller (--dont-use-soft-clipped-bases true, --standard-min-confidence-threshold-for-calling 20) and combined across the 12 samples with GenotypeGVCFs. We retained only biallelic SNPs that passed the following hard filters: QD ≥ 2.0, FS ≤ 30.0, SOR ≤ 3.0, MQ ≥ 40.0, MQRankSum ≥ –12.5, ReadPosRankSum ≥ –8.0, depth per site ≥ 10, and genotype quality ≥ 20. After filtering, 71,934 high-quality SNPs remained and were functionally annotated with snpEff v5.2f ^69^.

### Genomic substrates of overyielding and transgressive overyielding in growth

To detect loci where local genetic diversity contributes to diversity effects, we performed genome-wide higher-level association study (GHAS) ^18^ based on nucleotide diversity (*π*) computed for each 5-kb non-overlapping window for a given strain pair. For each window, the overyielding index was regressed against local *π* using linear models with the pairwise genetic relationship matrix (GRM) as a covariate. For transgressive overyielding, which exhibited zero-inflated and right-skewed distributions, we used generalized linear models assuming a Tweedie distribution (variance power = 1.5, log link) and again included the GRM to account for background genome-wide similarity.

Significant loci were identified based on *P*-values. Quantile–quantile plots and genomic inflation factors (*λ*_GC_) were used to assess the reliability of association results. For each phenotype, the top 50 loci were selected and defined as ±25-kb windows around the peak signal. Genes overlapping these intervals were extracted from the genome annotation file. Transcript IDs were mapped to orthologous gene identifiers using eggNOG-mapper annotations. These orthologs were linked to Gene Ontology (GO) terms, and GO enrichment analysis was conducted using Fisher’s exact test with Benjamini–Hochberg correction. GO terms were filtered for specificity by excluding overly broad terms (depth < 6 in the GO hierarchy). To identify protein-altering mutations within candidate genes, we interrogated the SnpEff-annotated VCF file. For transcripts located within ±25 kb of the top 50 loci identified by GHA analysis for each overyielding or transgressive overyielding metric, protein-altering variants classified as missense_variant, frameshift_variant, or stop_gained were extracted. For each variant, we tested whether individual phenotypes differed between genotypes and whether the corresponding overyielding or transgressive overyielding effect differed between strain pairs carrying the same versus different genotypes. Missense variants showing significant effects in both analyses were retained for subsequent PROVEAN analysis (Table S3). PROVEAN v1.1.5 ^70^ was used to estimate the intolerance for these non-synonymous substitutions, based on the evolutionary conservation and the chemical properties of the exchanged residues. Variants with PROVEAN scores ≤ −2.5 were considered deleterious and retained as candidates for potential functional divergence. We further predicted the three-dimensional protein structure using AlphaFold 3 ^71^. The resulting structural models were inspected to assess the spatial context of amino acid substitutions of interest. These variants were mapped onto the predicted structure and visualized using PyMOL (DeLanoScientific, San Carlos, CA), enabling examination of their positions relative to key structural elements and putative functional regions.

### Statistical analyses

All statistical analyses were done in *R* 4.4.1. The differences among strains in each trait were examined using analysis of variance (ANOVA), with strain as an explanatory variable and the phenotypes of each trait as response variables. To compare the population parameters (*r*, *K* and PC1) among strains and strain combinations, a linear mixed-effects model was performed using lmer functions in lme4 package. In this model, each population parameter was response variables, strain (or strain combinations) was a fixed effect, and medium condition was random effect. When calculating the value of overyielding and transgressive overyielding, I estimated the mean values population parameters for each strain and strain combination while accounting for medium conditions as random effects. First, a linear mixed-effects model was fitted using the lme4 package with strain (or strain combinations) as a fixed effect and medium condition as random effects. Then, the *emmeans* function in *emmeans* package was applied to obtain the estimated marginal means for each strain or strain combination, considering the specified random effects.

## Acknowledgments

We thank Todd P. Michael for providing the *Spirodela polyrhiza* 9512 genome assembly and annotation.

## Data availability

The datasets generated and analysed during this study, including phenotypic measurements, population growth data and source data underlying the figures, are available in the Figshare repository at 10.6084/m9.figshare.33878140. The raw RNA-Seq reads have been deposited in the NCBI Sequence Read Archive (SRA) under BioProject accession no. PRJNA1530017.

## Funding

This work was supported by JSPS KAKENHI Grant Number JP24K21984, research grants from the Asahi Glass Foundation and the Inamori Foundation, and the JST FOREST Program Grant Number JPMJFR253J, all awarded to Yuma Takahashi.

## Author contributions

Y.T. conceived the study. K.N., D.X.S. and Y.T. developed the methodology. K.N. performed the investigation. K.N., D.X.S. and Y.T. conducted the formal analyses. K.N. and Y.T. developed the software. Y.T. curated the data, prepared the visualizations, acquired funding, supervised the study and administered the project. K.N., D.X.S. and Y.T. wrote the original draft. All authors reviewed and edited the manuscript.

## Competing interests

The authors declare no competing interests.

## Supplemental Information

**Table S1.**
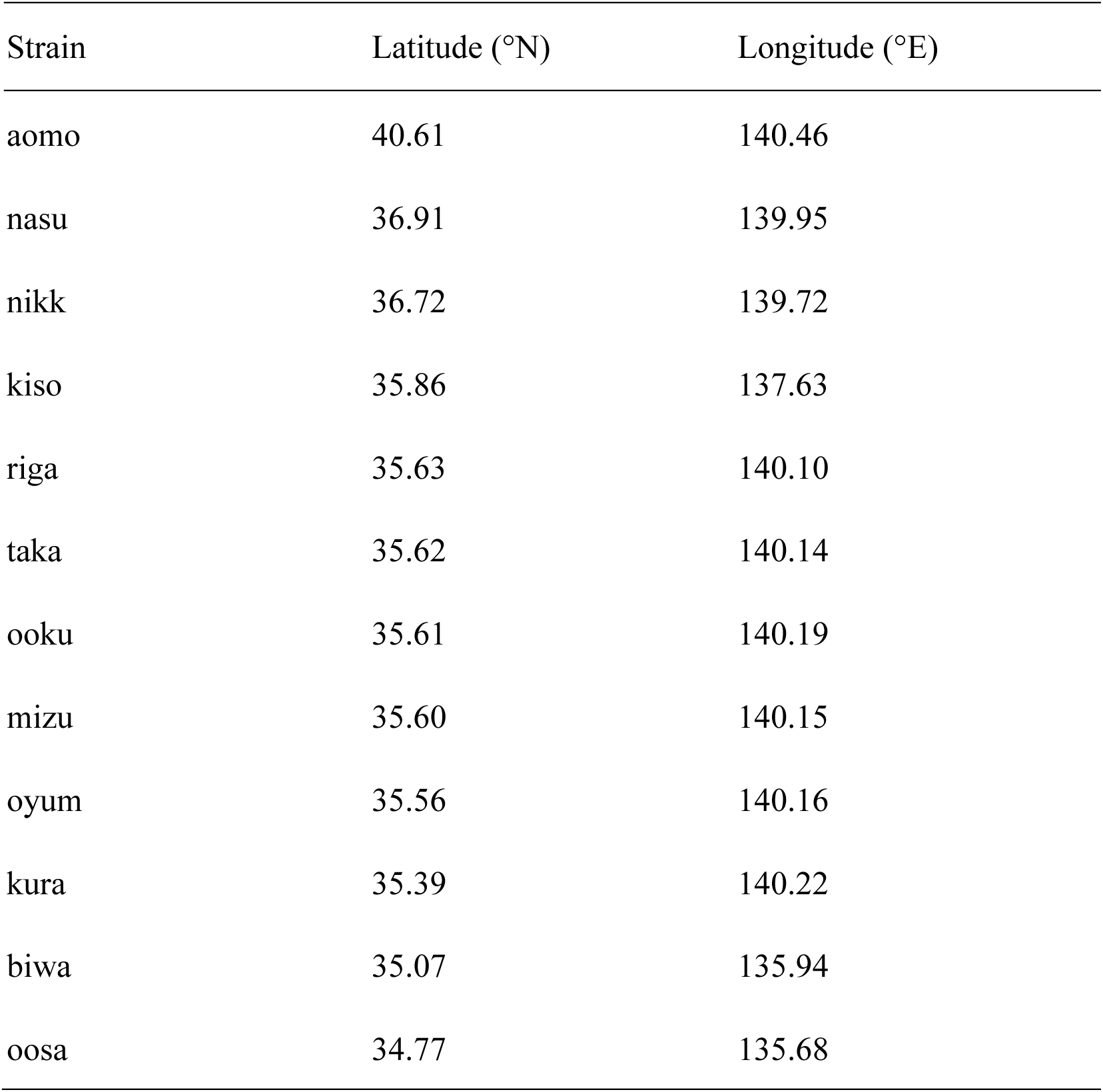
Strains and sampling sites.

| Strain | Latitude (°N) | Longitude (°E) |
| --- | --- | --- |
| aomo | 40.61 | 140.46 |
| nasu | 36.91 | 139.95 |
| nikk | 36.72 | 139.72 |
| kiso | 35.86 | 137.63 |
| riga | 35.63 | 140.10 |
| taka | 35.62 | 140.14 |
| ooku | 35.61 | 140.19 |
| mizu | 35.60 | 140.15 |
| oyum | 35.56 | 140.16 |
| kura | 35.39 | 140.22 |
| biwa | 35.07 | 135.94 |
| oosa | 34.77 | 135.68 |

**Table S1.** Information of sequenced samples and reads.

| Strain | Raw reads<br>(pairs) | Filtered<br>reads | % out<br>of raw<br>reads | Uniquely<br>mapped reads | % out of<br>filtered<br>reads | RIN |
| --- | --- | --- | --- | --- | --- | --- |
| aomo | 30,315,710 | 28,272,128 | 93.3 | 26,146,584 | 92.5 | 7.3 |
| biwa | 33,929,251 | 31,714,629 | 93.5 | 29,111,916 | 91.8 | 7.2 |
| kiso | 25,704,955 | 23,921,041 | 93.1 | 21,881,896 | 91.5 | 7.6 |
| kura | 22,413,818 | 20,759,676 | 92.6 | 18,610,593 | 89.6 | 7.5 |
| mizu | 27,267,377 | 25,210,691 | 92.5 | 22,046,653 | 87.4 | 7.9 |
| nasu | 32,851,449 | 30,610,151 | 93.2 | 27,897,649 | 91.1 | 7.7 |
| nikk | 28,383,770 | 26,439,362 | 93.1 | 23,363,287 | 88.4 | 8 |
| ooku | 24,337,483 | 22,687,210 | 93.2 | 20,460,900 | 90.2 | 7.7 |
| oosa | 28,129,720 | 26,258,414 | 93.3 | 24,037,473 | 91.5 | 7.5 |
| oyum | 29,326,850 | 27,400,167 | 93.4 | 24,980,361 | 91.2 | 7.8 |
| riga | 28,926,511 | 26,986,678 | 93.3 | 24,394,616 | 90.4 | 8.2 |
| taka | 29,686,715 | 27,775,077 | 93.6 | 25,190,726 | 90.7 | 8.1 |

**Table S2.**
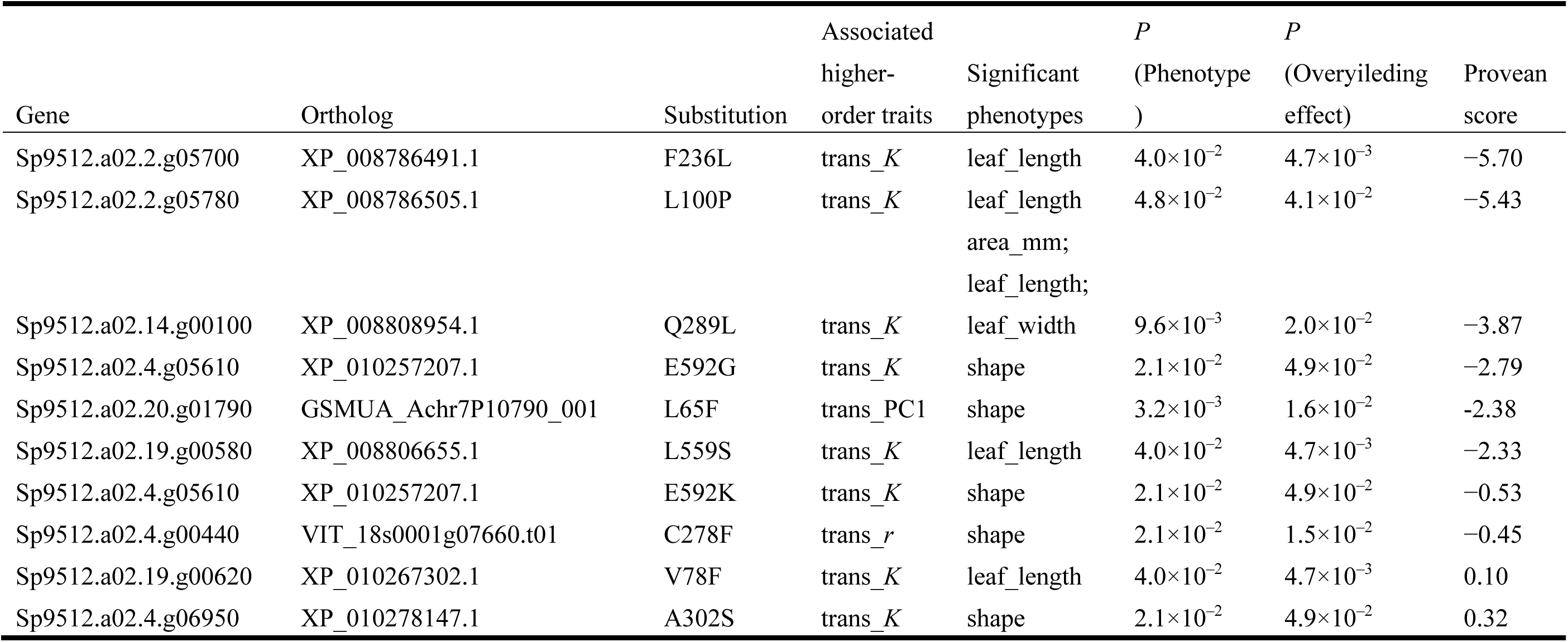
Candidate amino acid substitutions identified by GHA and phenotypic association analyses. PROVEAN scores ≤ −2.5 indicate substitutions predicted to have deleterious effects.

**Figure S1.**
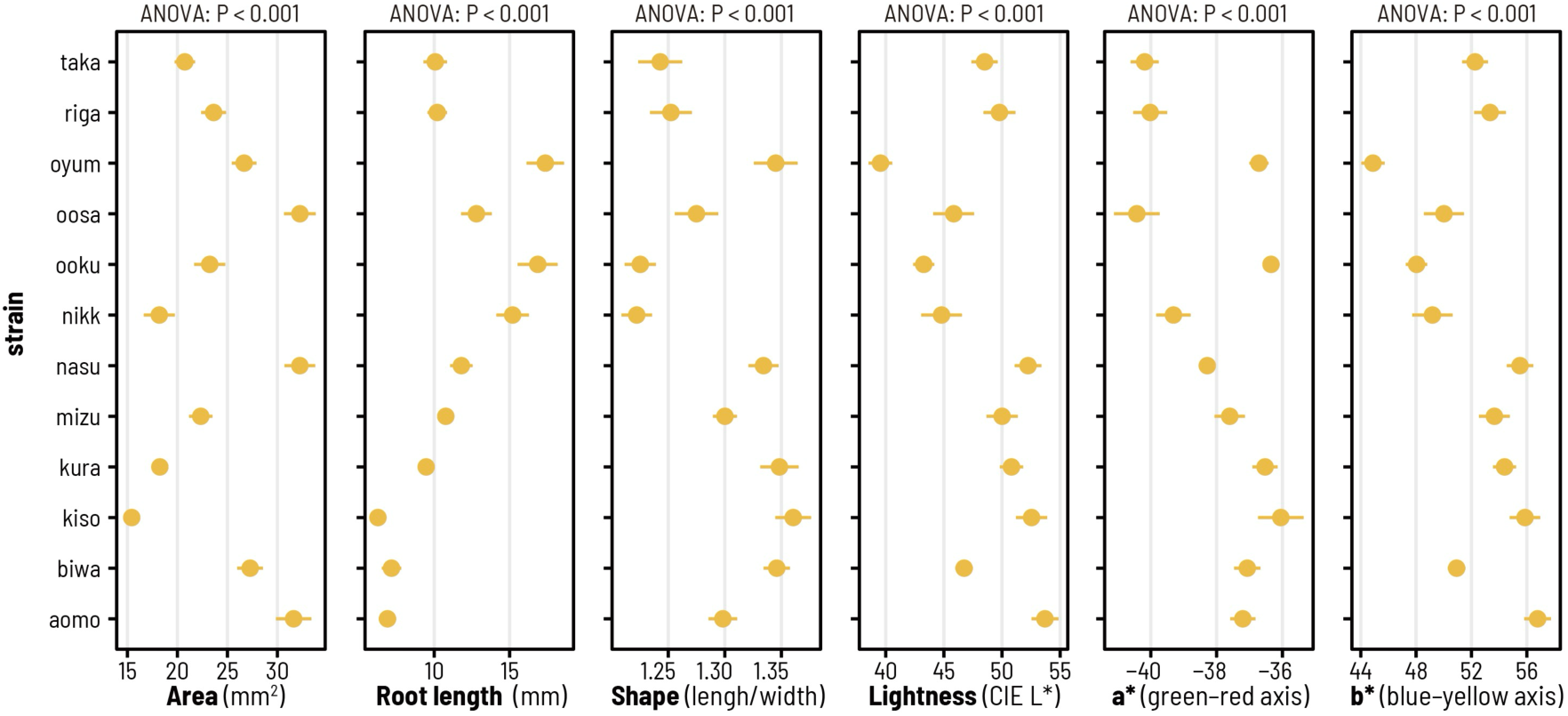
Among-strain variation in three morphological traits and three color parameters. Points indicate strain means, and horizontal bars indicate ± SE. All six traits differed significantly among strains (ANOVA, P < 0.001 for each trait).

**Figure S2.**
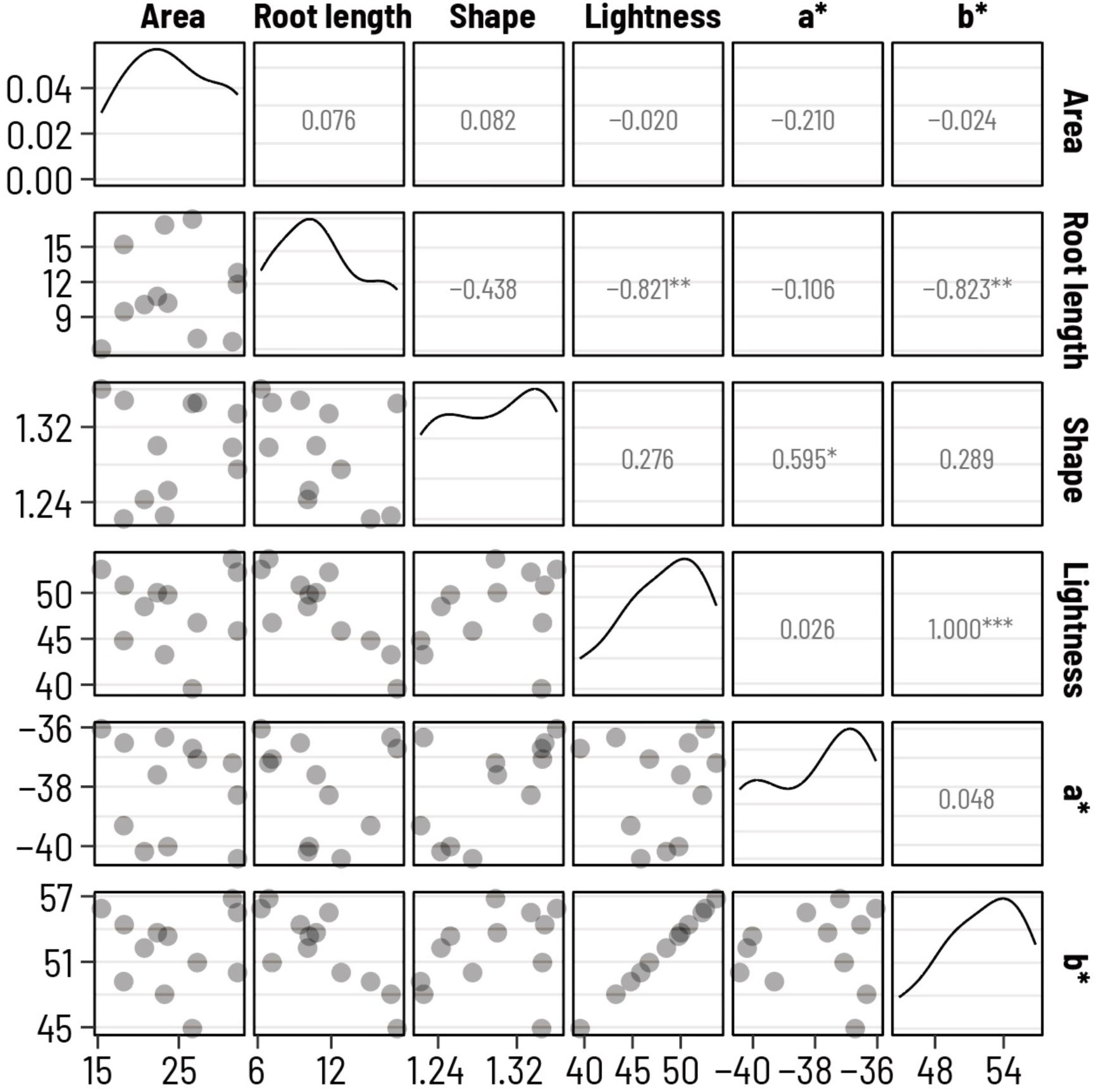
Correlations among three morphological traits and three color traits across strains. Scatterplots show pairwise relationships among frond area, root length, frond shape, lightness (L*), a* value, and b* value. Each point represents a strain. Values in the upper panels indicate correlation coefficients, with asterisks indicating statistical significance.

**Figure S3.**
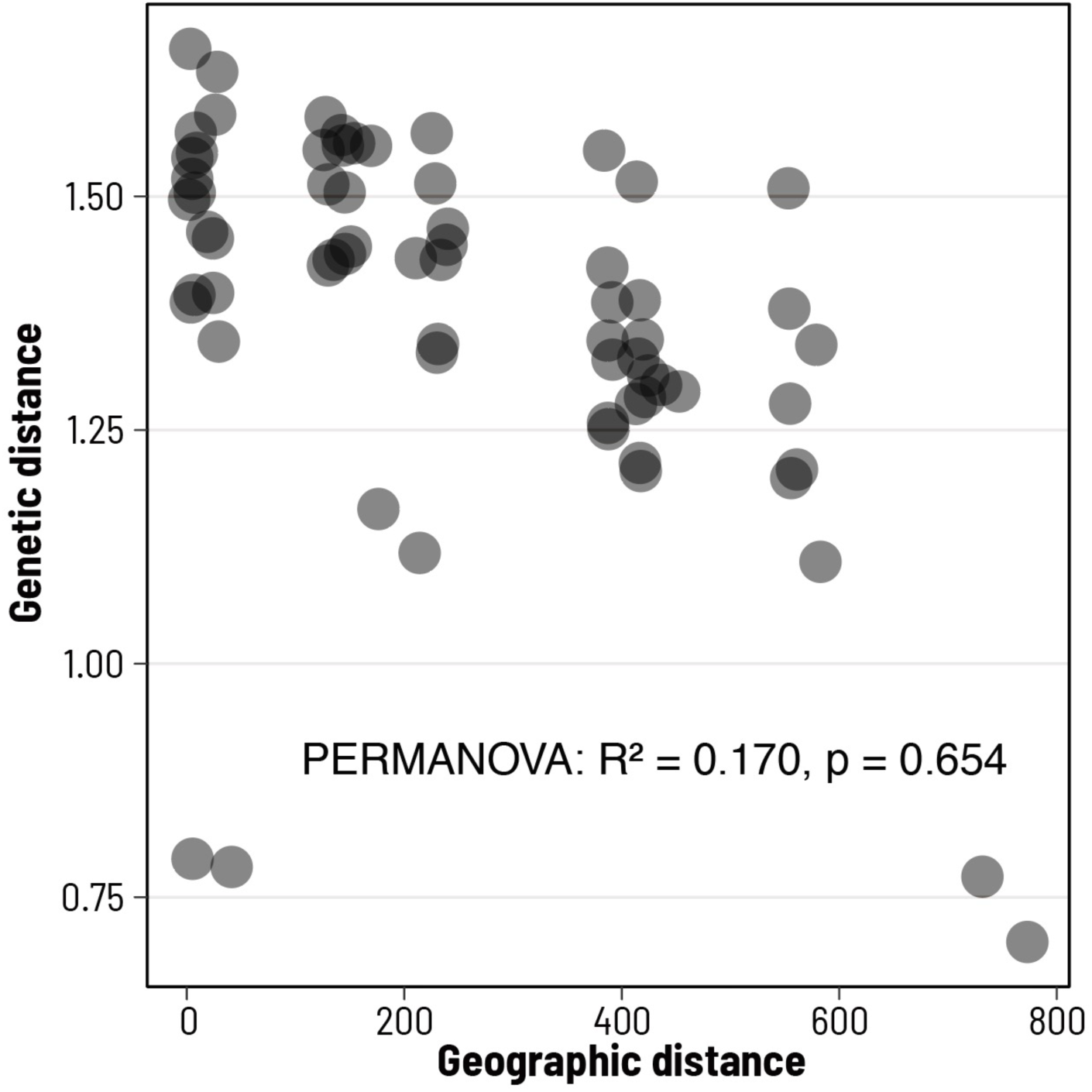
Relationship between geographic and genetic distances among strains. Each point represents a pairwise comparison between strains. Geographic distance was not significantly associated with genetic distance (PERMANOVA: R² = 0.170, P = 0.654).

**Figure S4.**
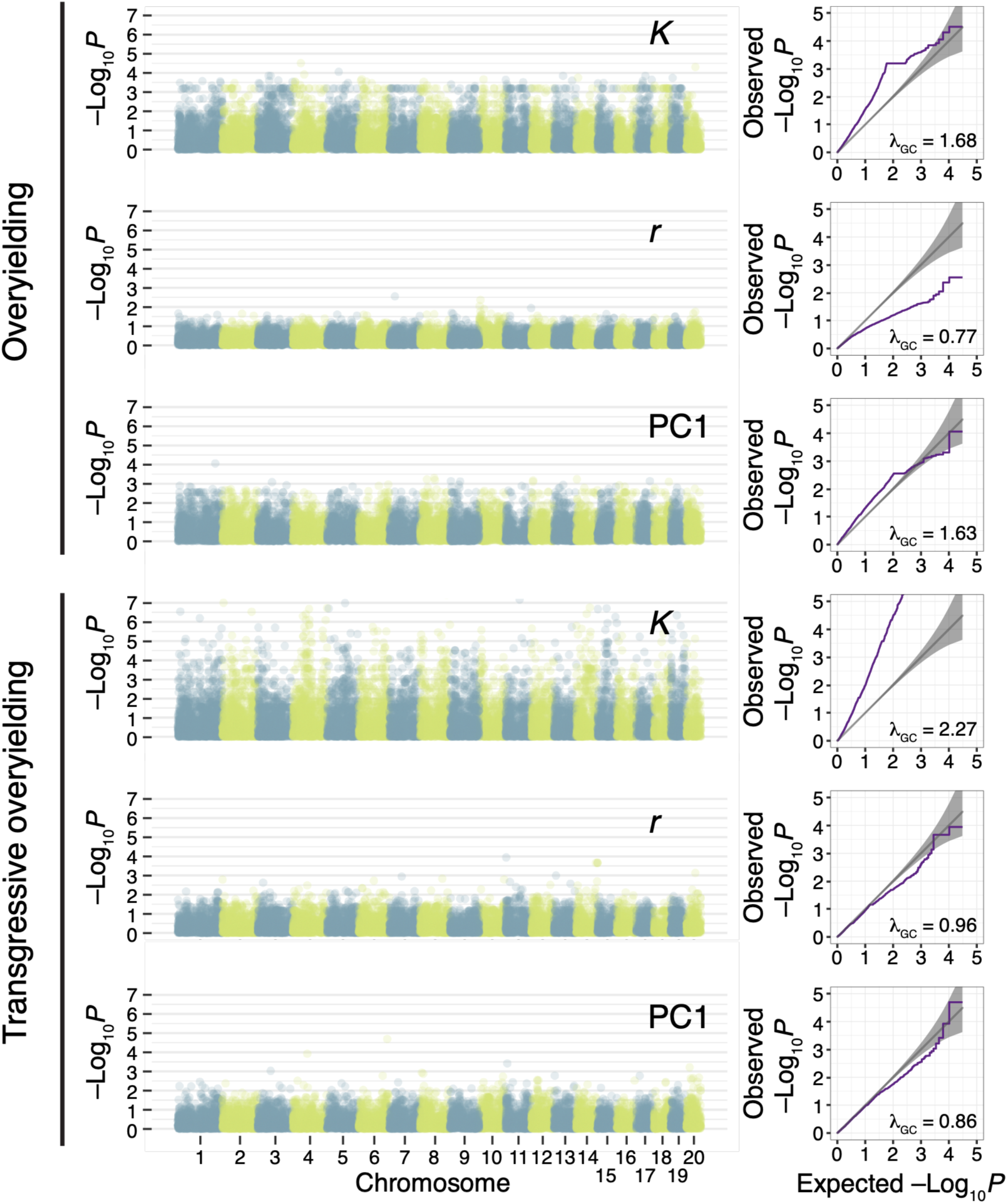
Genome-wide association profiles for overyielding and transgressive overyielding. Manhattan plots show genome-wide associations for carrying capacity (*K*), growth rate (*r*), and PC1 for overyielding (top) and transgressive overyielding (bottom). Alternating colors indicate chromosomes, and the y-axis shows −log_10_(*P*). Quantile–quantile (Q–Q) plots are shown to the right of each Manhattan plot, with the corresponding genomic inflation factor (λ_GC_).

**Figure S5.**
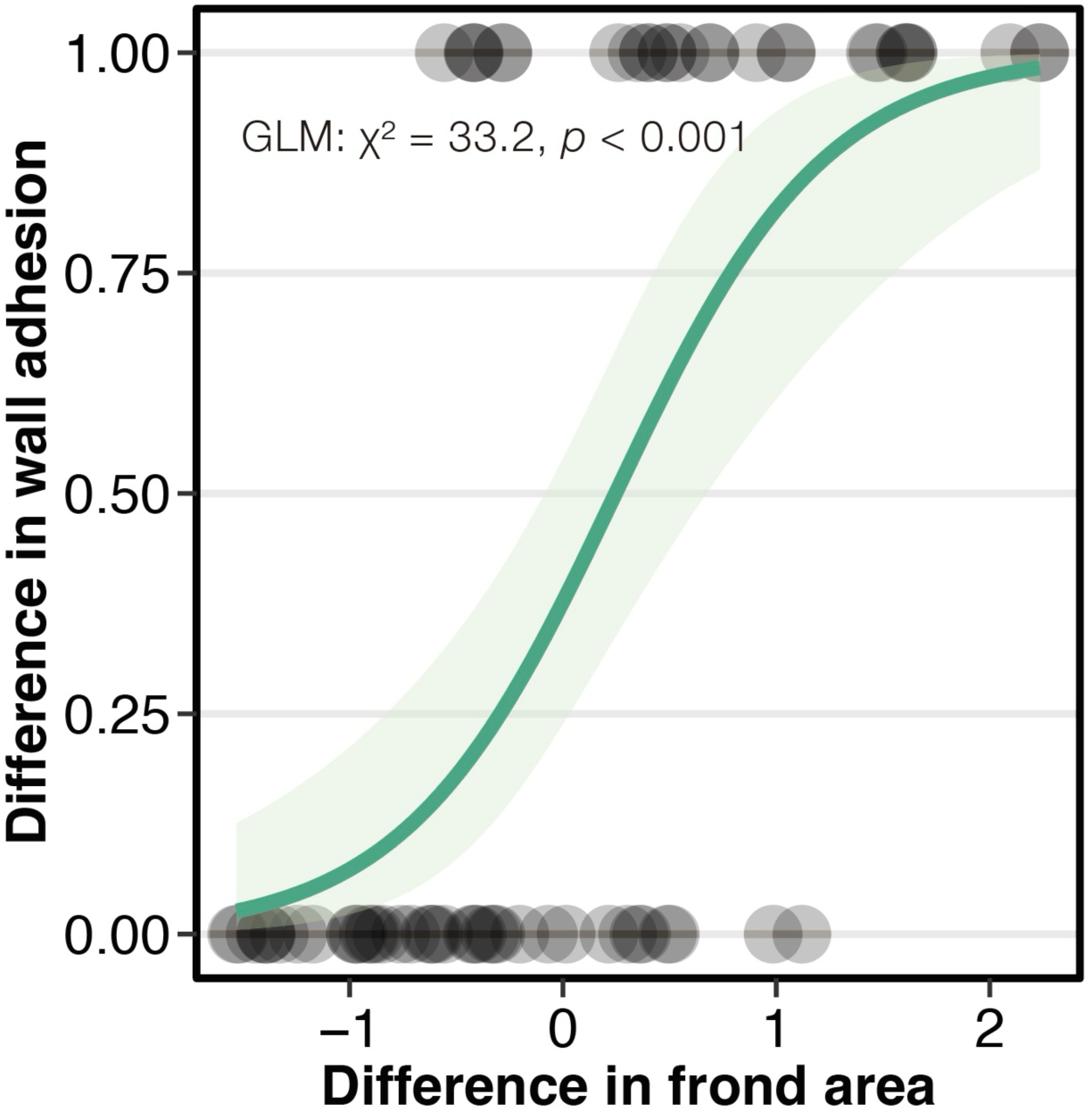
Relationship between differences in frond area and wall adhesion among strain pairs. Differences in wall adhesion were calculated by assigning strains that adhered to the wall a value of 1 and strains that did not a value of 0, and then taking the pairwise distance between strains. Pairs of strains with greater differences in frond area were more likely to differ in wall adhesion. Points represent strain pairs, and the solid line shows the fitted GLM with its confidence interval. The relationship was significant (GLM: χ² = 33.2, P < 0.001).

## Notes

### Competing Interest Statement

The authors have declared no competing interest.

